# Microbiota depletion promotes human rotavirus replication in an adult mouse model

**DOI:** 10.1101/2021.04.15.439998

**Authors:** Roberto Gozalbo-Rovira, Cristina Santiso-Bellón, Javier Buesa, Antonio Rubio del Campo, Susana Vila-Vicent, Carlos Muñoz, María J. Yebra, Vicente Monedero, Jesús Rodríguez-Díaz

**Author notes:** These two authors contributed equally (Roberto Gozalbo-Rovira, Cristina Santiso-Bellón). To whom correspondence should be addressed: Jesús Rodríguez-Díaz, Department of Microbiology, School of Medicine, University of Valencia, Avda. Blasco Ibáñez 17, 46010 Valencia, Spain, Vicente Monedero, Department of Biotechnology, IATA-CSIC, Av. Agustín Escardino 7, 46980 Paterna, Valencia, Spain, (Ext. 2006).

## Abstract

The study of human rotavirus (RV) infectivity *in vivo* has been limited by the lack of small animal models able to efficiently replicate the principal human RV genotypes. In recent years, intestinal microbiota-virus-host interaction has emerged as a key factor in mediating enteric virus pathogenicity. With the aim of developing an adult mouse infection model for RV we performed faecal microbiota transplant (FMT) with healthy infants as donors in antibiotic-treated mice. Contrarily to control mice, in the FMT group, but also in antibiotic-treated mice without FMT, challenge with the human RV G1P[8] genotype, Wa strain (RV_wa_), resulted in viral shedding in the faeces for 6 days. RV titres in faeces were also significantly higher in antibiotic-treated animals with or without FMT. This excluded the hypothesis that donor’s microbiota promoted infection. Antibiotic treatment followed by self-FMT resulted in incomplete re-establishment of mouse microbiota which partially restored suppression of RV_wa_ infection. Microbial composition analysis revealed profound changes in the intestinal microbiota of antibiotic-treated animals, whereas some bacterial groups, including members of *Lactobacillus, Bilophila, Mucispirillum* and *Oscillospira*, reappeared after self-FMT. In antibiotic-treated and FMT animals, differences were observed in gene expression of immune mediators such as IL10, TNF-α and IFNγ and the fucosyltransferase FUT2, responsible for H-type antigen synthesis in the small intestine. Collectively, our results suggest that antibiotic-induced microbiota depletion eradicates the microbial taxa that restrict human RV infectivity in mice. Viral permissiveness could involve changes in the innate immune system at the small intestine and alterations in the bacteria population that potentially interact with RV, creating a favourable environment for human RV replication in mice.

## Introduction

Diarrheal disease is the second leading cause of death worldwide in children under five years of age, accounting for around 525,000 deaths each year. Rotavirus (RV) is among the predominant causes of non-bacterial acute gastroenteritis in infant and young children, with an estimated 150,000 deaths per year, mostly in developing countries [1]. The gut is a very complex ecosystem, with multiple interactions between the host immune system, glycobiology, resident microbiota and viruses responsible for gastroenteritis [2–5]. A link between human RV infection and intestinal bacterial populations has been revealed from analysis of the microbiota in population groups displaying different vaccine take after inoculation with RV live vaccines [6,7]. However, the role of commensal microbiota in RV infectivity is still controversial. Some evidence points towards a positive effect, showing reduced RV infectivity in germ-free or antibiotic-treated mice [8]. Contrarily, some microorganisms, including probiotic bacteria, directly interact with RV and potentially block their binding to epithelial receptors, counteracting infection [9].

RV attach to glycans of the histo-blood group antigen (HBGA) expressed in the gastrointestinal tract. In humans, fucosylation of O-linked HBGA at this location is controlled by *FUT2* and *FUT3* gene expression, and it has been demonstrated that the glycosylation patterns of those receptors could be altered by the action of specific commensal bacteria [10]. The main human RV genotype (G1P[8]) recognizes fucosylated HBGAs such as H-type 1 [11], synthesized by FUT2 activity, and Lewis b antigens [12], synthesized by FUT3, but these viruses show inefficient spread or no replication at all in the mouse model [13].

With the aim of obtaining a versatile animal model for human RV infection we tested whether engraftment of infant intestinal microbiota would permit G1P[8] RV infection in the mouse model. We showed that a simple microbial ablation through antibiotic use was sufficient to promote human RV infection in mice, highlighting a role for intestinal microbiota in supressing these RVs in the murine model.

## Results

### Mice with depleted microbiota shed Wa rotavirus for 6 days

We reasoned that in the intestinal niche of infants where RV develop, the resident microbiota determines infectivity. Then, we asked whether infection of RV_wa_ (G1P[8]) in the mouse model could be improved by replacing its intestinal microbiota with the microbiota of infants by FMT using antibiotic (Ab)-mediated microbiota ablation. In order to answer this, C57BL/6J mice were treated with an antibiotic cocktail through drinking water for three weeks. Bacterial counts in faecal pellets decreased from 6.3-9.7 x 10^10^ CFU/g to no detectable counts (plating 10^−1^ dilutions), whereas they remained stable in the non-treated control (4.9 x 10^10^ CFU/g), indicating an efficient depletion of the culturable gut bacteria. Mice were then subject to FMT and orally dosed with the Wa strain. The virus could be detected in the faeces of control animals with a peak at one day post-infection (dpi; 9.1 x 10^9^ genome copy equivalent (GCE)/g faeces), which rapidly dropped, with RV being under the detection limit at dpi 4 (Fig. 1). By the contrary, compared to control animals, at dpi 1 viral shedding in Ab-treated mice and in mice subject to human FMT (hFMT) was 23- and 16-fold higher (2.1 x 10^11^ GCE/g and 1.5 x 10^11^ GCE/g, respectively) and remained until dpi 5-6, following the same evolution pattern and with values higher than 10^9^ GCE/g. The lower levels of RV detected in control animals and its faster clearance were indicative of an inefficient replication, in contrast to the prolonged viral shedding in Ab-treated mice and in the hFMT group. In mice receiving a self-FMT, partial restoration of the restrictive replication of RV_wa_ was evidenced, and virus shedding was detected at levels similar to control at dpi 1 and only increased 10-fold with respect to control at dpi 2 and 3, with no detectable viral shedding at dpi 4 (Fig. 1).

**Figure 1.**
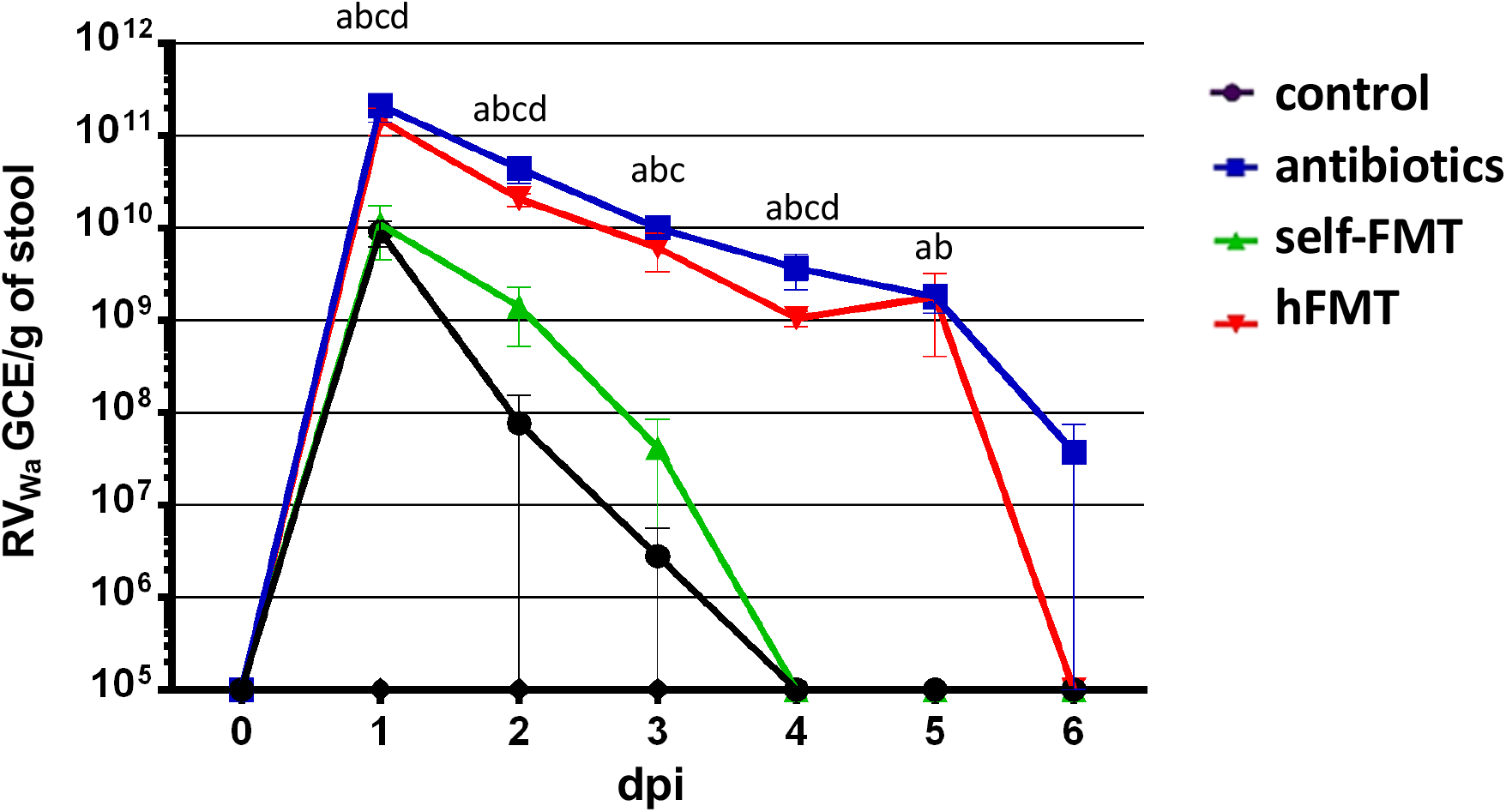
Rotavirus Wa shedding in mice stool. GCE/g of stool are presented for control mice, Ab-treated mice and mice subject to hFMT and self-FMT, respectively. The mice were followed for 6 days and euthanized on day 7. The letters indicate statistically significant differences (*p*<0.05) in the GCE between the groups (a, Ab vs control; b, Ab vs self-FMT; c, hFMT vs control; d, hFMT vs self-FMT). The error bars are standard deviations (n=5). The limit of detection was 10^5^ GCE/g of stool. Ab, antibiotic; hFMT, human fecal microbiota transplantation; self-FMT, self-fecal microbiota transplantation.

### Impact of faecal microbiota transplant on mouse microbiota

The results of viral shedding pointed to microbiota eradication by Ab rather than hFMT as responsible for RV_wa_ infection permissiveness in mice. The microbiota composition in faeces prior to RV challenge was determined by 16S rDNA NGS, showing that bacterial richness and diversity was severely affected after the different treatments (Fig. 2A). Chao1 index was lowered in all groups compared to control (microbial richness; *p* = 2.0365e-12; F = 206.49, ANOVA) but was higher in self-FMT animals. Diversity (Shannon index) was also lower (*p* = 1.8988e-08; F= 57.299, ANOVA), but was higher in the hFMT group than the self-FMT group, indicating uneven microbial distribution after treatments. In both type of analyses Ab treatment resulted in lower a diversity. As expected, the three distinct treatments caused profound remodelling of the intestinal microbial profile and all groups could be differentiated in terms of overall composition (Fig. 2B and 2C). In mice treated with Ab, members of the phyla Firmicutes and Proteobacteria, belonging to the genus *Lactococcus* and unidentified γ-Proteobacteria (enterobacteria), respectively, accounted for most of the microbiota (Fig. 2B). The microbiota present in the pooled infant faeces was not completely engrafted in mice after hFMT (Supplementary Fig. S1), but these mice were characterized by elevated numbers of Bacteroidetes and presence of members of the genus *Megasphaera*, a bacterial taxon that was absent in the microbiota of untreated mice (Fig. 3A). Genera such as *Bifidobacterium, Adlercreutzia, Ruminococcus, Coprococcus, Turicibacter, Odoribacter* and *Allobaculum* were completely depleted after Ab treatment and they were not replenished by any FMT, including self-FMT (Fig. 3B). Although microbial engraftment was not completely successful in the self-FMT group (differences in α and β diversity were obvious compared to control animals), bacteria belonging to *Oscillospira, Lactobacillus, Mucispirillum* and *Bilophila* were partially restored in mice via self-FMT (Fig. 3C). In addition, in self-FMT animals proportions of *Akkermansia* and *Sutterella* taxons were increased compared to control mice (Fig. 3D). In summary, low microbial richness and ablation of most representative taxa of mice microbiota was concomitant with higher mice permissiveness to RV_wa_ infection.

**Figure 2.**
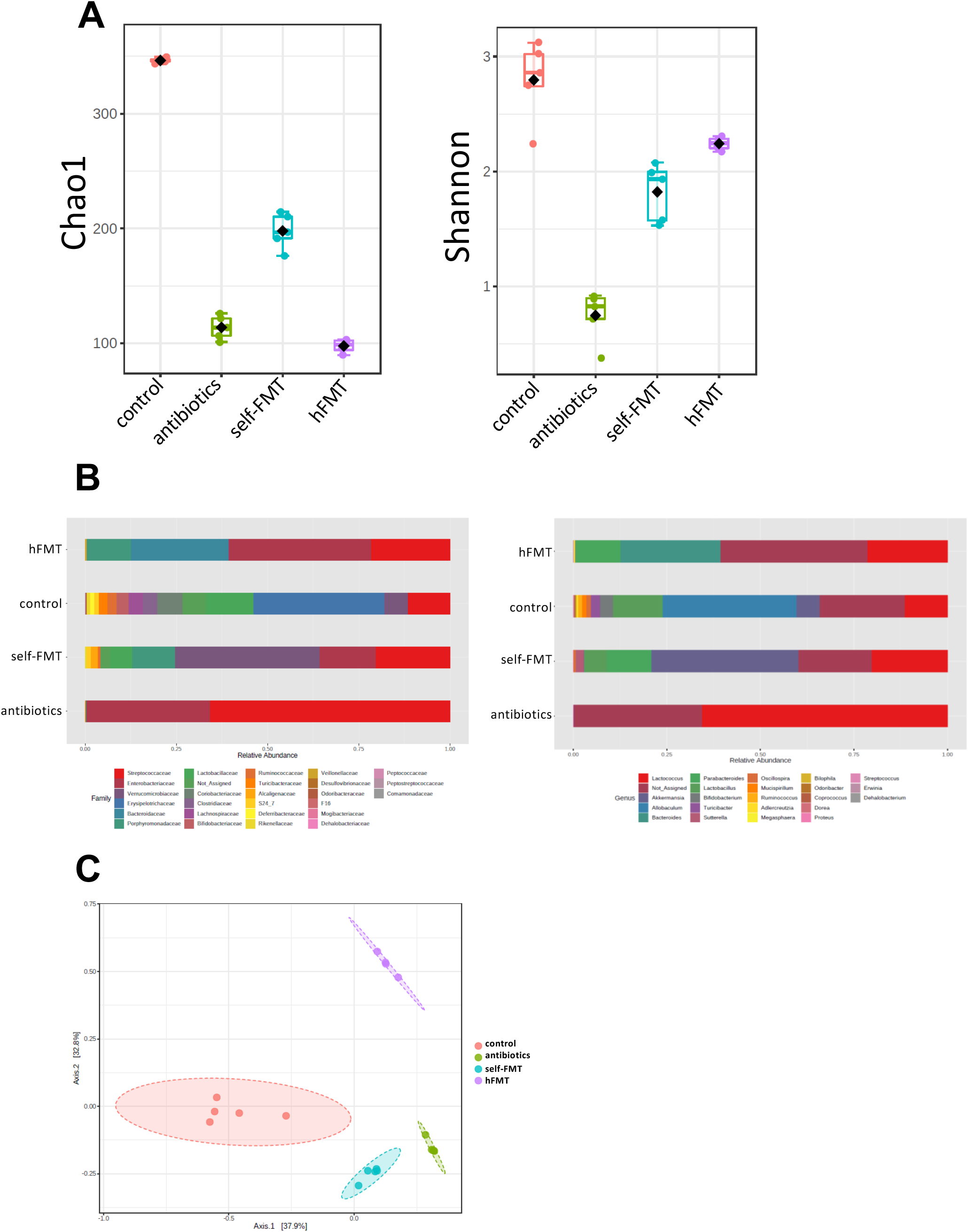
Changes in intestinal bacterial composition in the mice groups. **(A)** Microbial α-diversity. Microbial richness (Chao1) and diversity (Shannon) indexes are shown. **(B)** Relative abundance of bacterial taxa (family and genus levels) found in the faeces of the different mice groups. **(C)** Differences in microbial global composition after the different treatments. A principal coordinate analysis (PCoA) of the Bray-Curtis dissimilarity indexes of samples is shown (PERMANOVA F-value = 36.77; R^2^ = 0.8803; *p*<0.001).

**Figure 3.**
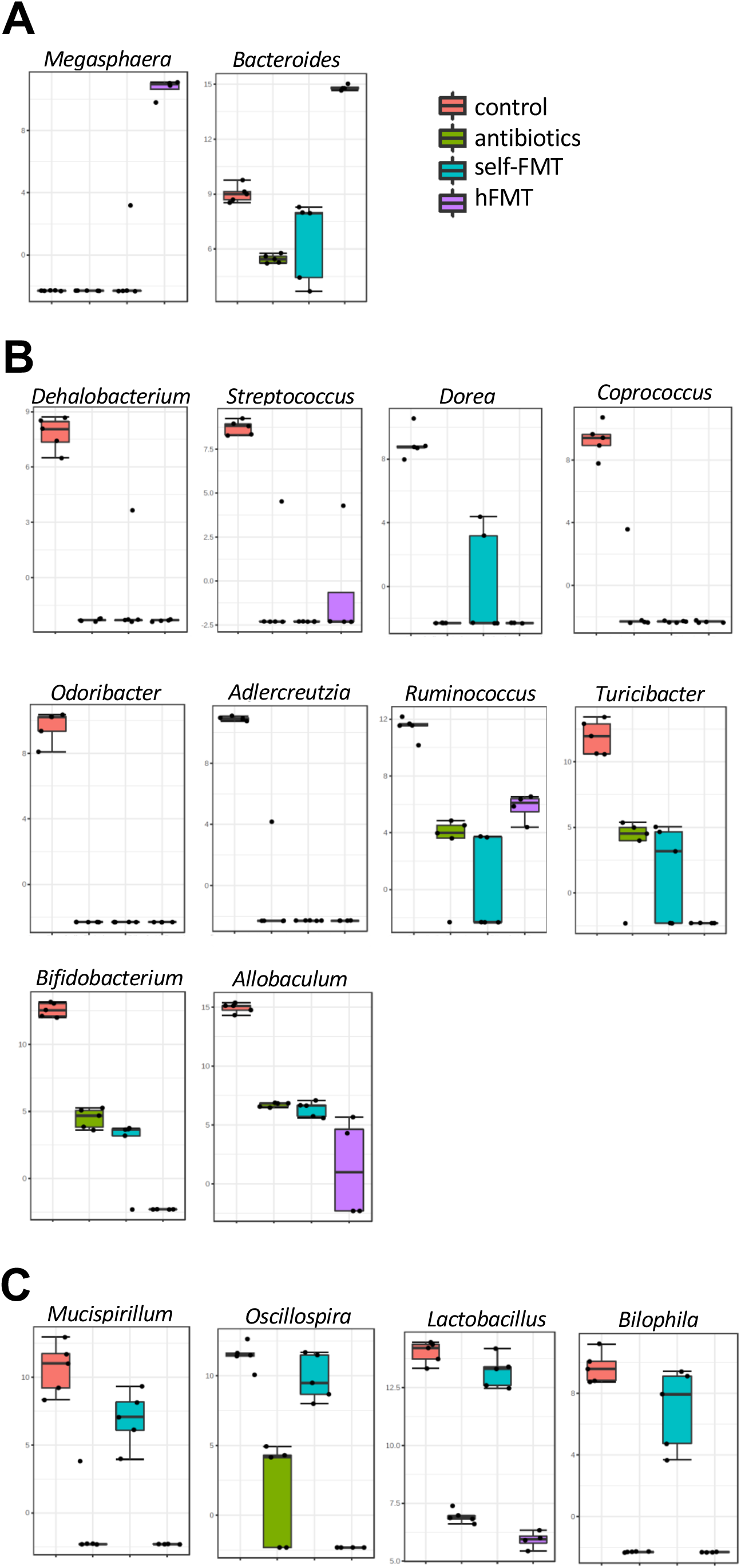
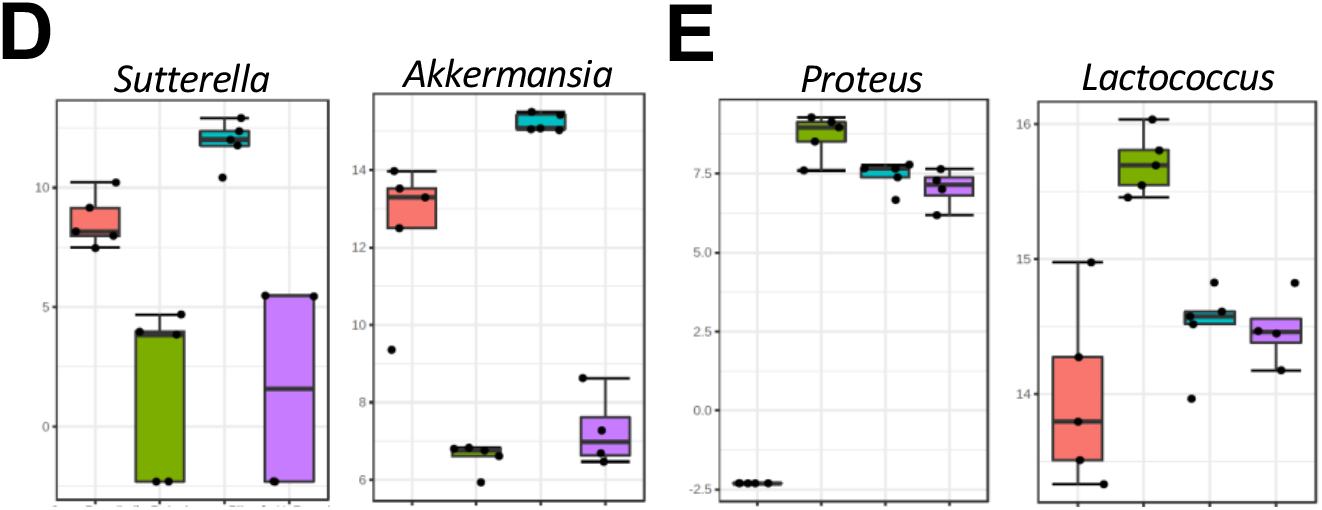
Bacterial genera showing differences between treatment groups. A LEfSe analysis was performed to detect bacterial markers characteristic of each treatment condition. The different box-plots represent abundance (log transformation of data normalized by total-sum scaling x 10^7^) of bacterial genera showing LDA scores >4. **(A)** Bacteria augmented after hFMT; **(B)** Genera ablated or diminished by Ab treatment; **(C)** Bacteria that were partially restored by self-FMT; **(D)** Bacteria augmented after self-FMT; **(E)**. Bacteria increased in Ab-group. Red, control animals; green, antibiotics-treated animals; blue, self-FMT; magenta, hFMT.

### Microbiota depletion with antibiotics affects immune and epithelial glycosylation gene expression in infected mouse gut

Gene expression levels in a panel of cytokines and other innate immune system mediators including IL1β, IL4, IL6, CXCL15, IL10, IL12, IL13, IFNγ, TNFα and TLR2 were studied by RT-qPCR in the small intestine of mice at 7 dpi (Fig. 4). *IL6, IL12* and *IL13* expression levels did not differ between experimental groups and control animals, whereas a downregulation effect was generally observed in the remaining tested genes. *IL1β* and *CXCL15* messenger levels were lower in the Ab and hFMT groups. Expression of *IL10, TLR2* and *TNFα* was also reduced in all groups compared to control. Finally, *IFNγ* expression was diminished in mice subject to both FMT treatments. Expression of the *FUT2* gene, whose product is involved in α1,2-fucosylation during the synthesis of mucosal H type-1 antigen (fucosyl-α1,2-galactopyranosyl-β1,3-*N*-acetyl-glucopyranoside), one of the P[8] RV adhesin receptors, was downregulated in self-FMT animals.

**Figure 4.**
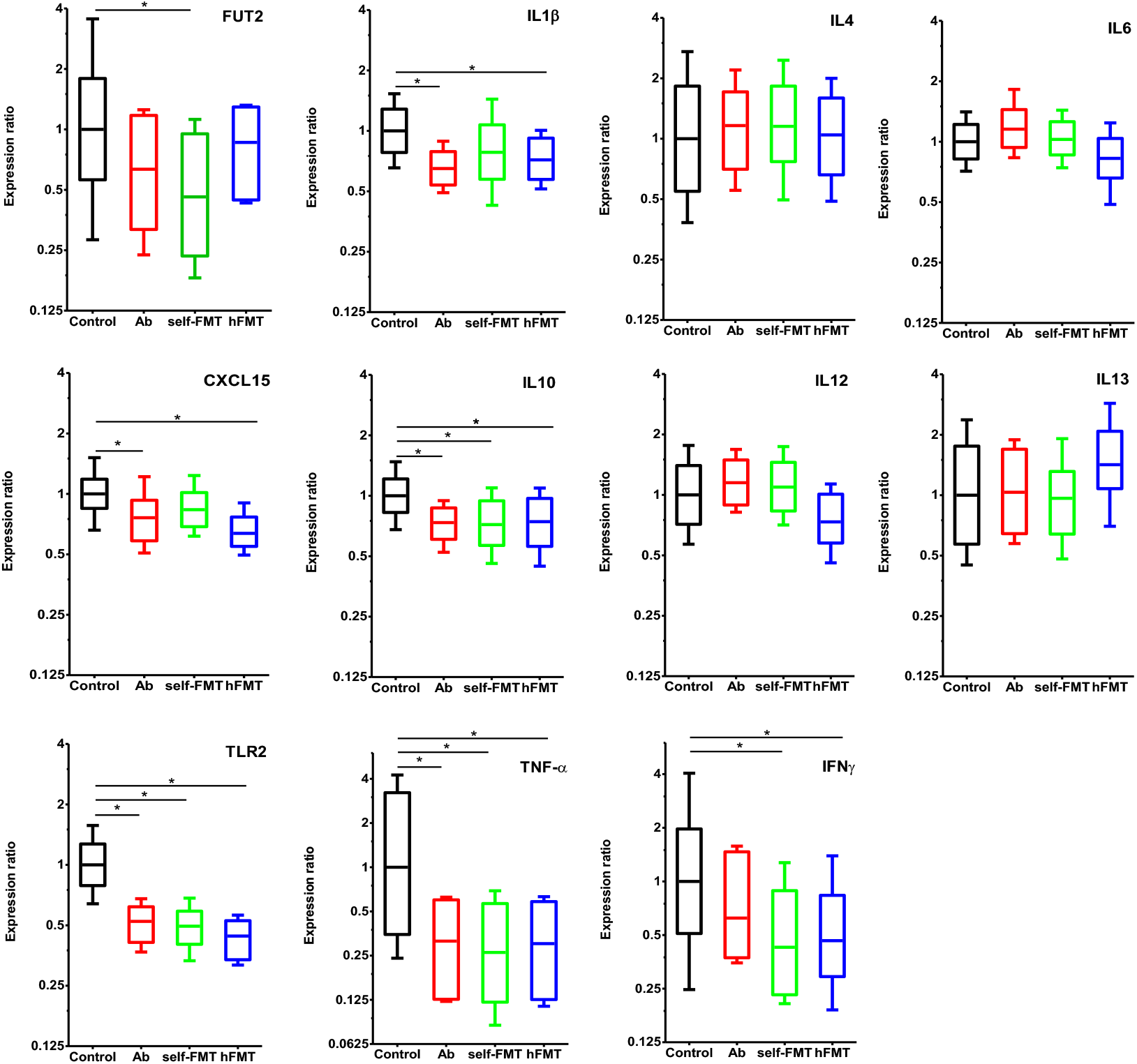
Differences in gene expression in the small intestine of infected mice. Expression of immune system mediators (IL1β, IL4, IL6, CXCL15, IL10, IL12, IL13, IFNγ, TNFα and TLR2) and fucosyltransferase 2 (FUT2) was determined by RT-qPCR in mice from the different groups at 7 dpi (n=5). The level of significance is indicated for control mice (*p*<0.05).

## Discussion

Gut microbiota has emerged as a pivotal player in enteric virus-host interactions, and has been shown or hypothesized to have positive as well as negative effects on viral infectivity, mediated by different mechanisms [2–5]. Given the lack of an efficient mouse model to replicate relevant human RV strains [13], we studied whether mice could be infected with RV_wa_ by infant microbiota engraftment. This RV strain has been used as a model for human RV studies and has special relevance because it carries the P[8] VP4 genotype, which is responsible for around 90% of infections [14]. An available *in vivo* model of replication would therefore be desirable, yet its infectivity in small animal models is not satisfactory.

In our study, RV_wa_ infection could not be linked to hFMT, ruling out the possibility that human intestinal microbiota mediate RV_wa_ infectivity in mice. Nonetheless, we show that independently of subsequent hFMT, simple gut bacteria depletion through antibiotic treatment dramatically increased the replication capacity of the RV_wa_ strain in this host. Our data suggest that endogenous mouse microbiota restrict RV_wa_ replication, which is in opposition to results obtained with the murine RV EC strain, where Ab treatment had negative effects on viral entry, delaying and decreasing infectivity in both adult (40% reduction in viral shedding) and neonatal (34% reduction in diarrhoea incidence) murine models [8]. However, recent research also points to a protective role for the microbiota during murine RV (strain EDIM) infection, by demonstrating high susceptibility to RV infection in Ab-treated and germ-free animals and highlighting a role of microbiota-induced IL22 expression as an antiviral mediator [15].

The importance of the microbiota in establishing an appropriate anti-viral immune response has been noted [16,17]. Exacerbated systemic infection by several enteric and non-enteric viruses was observed in Ab-treated mice, including vesicular stomatitis and influenza virus [18], murine gammaherpesvirus [19], respiratory syncytial virus [20], encephalomyocarditis virus [21], and West Nile, Dengue and Zika viruses [22]. In some cases, microbiota depletion has been linked to a defective innate immune response characterized by low levels of type I IFN expression (IFNβ), which hampered the ability to mount an effective antiviral macrophage response [18,20]. In our experiments, antibiotic treatment generally resulted in reduced expression of inflammatory mediators in the small intestine, although no *IFNβ* expression could be detected in the whole tissue (data not shown). We found that *IL10* and *TNFα* displayed reduced expression in microbiota-depleted animals and that *IFNγ* expression was also lower in FMT animals. These are important mediators of immune response to infections, including RV, in which studies have previously reported increased expression after infection in a mouse model [23,24]. Accordingly, although germ-free mice show delayed infection with EC RV, viral clearance in this model was found to last substantially longer, probably reflecting an immature intestinal immune system [8]. Therefore, reduced IL10, TNFα and IFNγ are likely to be beneficial for viral replication. TLR2, whose expression was lowered in the Ab-treated groups, has been linked to the protective effect of lactobacilli towards RV infection in animal models [25,26]. We also studied the association between RV infection and *FUT2*. It was previously shown that specific members of the gut microbiota such as *Bacteroides thetaiotaomicron, Bacteroides fragilis* and segmented filamentous bacteria (SFB) are able to influence the pattern of mucosal glycosylation by inducing epithelial FUT2 expression [10,27], which encodes the enzyme responsible for H-type antigen synthesis, the target for binding of RV P[8] adhesin [11]. We showed that *FUT2* expression in the small intestine was only reduced in self-FMT animals. Lack of intestinal microbiota generally results in downregulation of FUT2 [27], and it is not known how diminished *FUT2* levels could affect RV_wa_ infection in mice, especially given that another entry mediator of P[8] RV, the Lewis b antigen [12], is not produced in this host. A recent report shows that virulent RV_wa_ (a strain serially passed in gnotobiotic pigs) infection in porcine enteroids is enhanced by the presence of H antigen [28], but the role of HBGA in infection in mice has yet to be determined.

The limited RV_wa_ infection found in self-FMT animals suggests that specific microbial taxa not implanted after hFMT are implicated in RV exclusion. Although self-FMT did not restore the original microbiota of mice, several bacterial taxa with a potential role in RV_wa_ restriction were partially restored: *Mucispirillum* is a strict anaerobe typically found in the murine intestinal tract, where it is intimately linked to the mucosal layer and participates in inflammatory processes [29] and is also involved in excluding several pathogenic bacteria [30]; *Oscillospira* is another anaerobe from the grastrointestinal tract, and although it has never been cultivated, evidences indicate its relevance to human health [31]; *Bilophila*, which comprises the single species *Biliophila wadsworthia*, has been linked to intestinal inflammation in mice [32]; and finally, lactobacilli are well-known probiotic bacteria that have been shown to diminish RV-induced diarrhoea in humans [33] and reduce RV infection in both *in vitro* and *in vivo* models [34,35].

A recent study showed that members of the murine intestinal microbiota such as SFB belonging to *Candidatus* Arthromitus are responsible for inhibiting murine RV EC strain infection [36]. Thus, Ab elimination of SFB could explain Ab-treated mice tolerance to RV_wa_ infection. However, SFB were not found in our microbiota analysis and the model where they were implicated in resistance to the murine RV strain EC used immunosuppressed Rag1-KO mice, which are defective in B and T lymphocytes. However, *Candidatus* Arthromitus was not able to stably colonize normal mice [36]. Nevertheless, the idea that some bacterial taxa, such as those re-established after self-FMT or others, could mediate RV_wa_ exclusion in mice is still attractive. Such a category of bacteria has been identified and the antiviral mechanisms behind them disclosed in some cases. As examples, oral dosing of *Blautia coccoides* in Ab-mice restored the capacity of macrophages to induce IFNβ and promoted protection against encephalomyocarditis virus systemic infection [21], and likewise, lipo-oligosaccharides from the outer membrane of *Bacteroides* and microbial metabolism-derived acetate were involved in triggering an IFNβ response that prevented vesicular stomatitis virus and influenza virus [18] and respiratory syncytial virus [20] infection in mice, respectively.

RV_wa_ permissiveness in Ab-treated mice could also derive from depletion of one or various of the many microbial taxa affected by this treatment but not restored by self-FMT. As an example, *Ruminococcus* practically disappeared after Ab administration. These bacteria were correlated with low anti-RV IgA titers in human saliva of healthy subjects [37], have recently been characterized as microorganisms physically interacting with RV in the stools of children suffering P[8] RV-induced diarrhoea, and reduced RV_wa_ infectivity in Caco-2 cell cultures [9]. *Bifidobacterium*, another relevant genus of probiotic bacteria with proven anti-RV properties [35,38] was also eradicated by antibiotics.

Alternative mechanisms by which bacteria could mediate RV protection include direct interaction of RV virions with bacteria, as hypothesized for SFB [36]. These interactions have been evidenced in *in vitro* experiments for a number of bacterial strains [23,39]. Species such as *Ruminococcus gauvreauii* [9] and others [40] express HBGA-like substances on their surface which are binding targets for RV. HBGAs act as virion stabilizers that promote bacteria-assisted binding and enhance infection in some enteric viruses, such as norovirus (NoV) and poliovirus [41–43] but can mediate virus sequestration on the bacterial surface, which could prevent virus from binding to target cells.

The mechanisms underlying RV_wa_ development in antibiotic-treated mice are as yet unknown, but our results suggest either that natural mouse resistance to RV_wa_ infection is triggered by indigenous mouse commensals that cannot be substituted by other human intestinal bacteria, or alternatively, that the equivalent human microorganisms cannot be implanted in the mouse by simple FMT. Further characterization of the factors mediating RV replication in Ab-treated mice is needed. Nonetheless, this *in vivo* infection model may constitute a valuable tool to investigate the biology of RV. Identifying the bacteria responsible for human RV restriction in mice will further understanding of the relationship established between RV and intestinal commensals.

## Methods

### Rotavirus stock preparation

Human RV Wa strain was produced as previously described [44] with modifications. Briefly, 10 MA104-confluent 150-cm^2^ flasks (approximately 1.5×10^7^cells/flask) were infected with Wa RV at a multiplicity of infection (MOI) of 0.1 and the flasks were kept at 37 °C for one week. After infection the virus was pelleted at 100,000 xXg for 2 h in a Himac R25ST-0507 rotor coupled to a Himac CR-30NX centrifuge. The pelleted virus was resuspended in TNC (20 mM Tris-HCl pH 8.0, 100 mM NaCl, 1mM CaCl_2_) and ultra-centrifuged in a sucrose gradient (30-70%) in TNC. The gradient was run in a SW41 rotor coupled to a Beckman L80 ultracentrifuge at 150.000 x *g* for two hours. The band containing RV was collected and the virus was finally recovered by pelleting in the SW41 rotor coupled to the Beckman L80 ultracentrifuge for 2 h at 150.000 x *g* and resuspended in TNC. After preparation the RV Wa stock was titrated by qPCR as previously described [45].

### Ethical statement

All experiments using mice were conducted at the Animal Production and Experimentation Service of the University of Valencia following national and international regulations. The procedures were approved by Ethics Committee for Animal Welfare and by the “Dirección General de Agricultura, Ganadería y Pesca” of the “Generalitat Valenciana”, file number 2018/VSC/PEA/0181. Use of human stool samples was approved by the Human Research Ethics Committee of the University of Valencia (registration number H154401046838) and written informed consent was obtained from a parent of each subject.

### Donor microbiota preparation

Stool samples from four healthy human infants between one and three months of age were collected and resuspended in a solution consisting of 80% 2X-concentrated brain-heart infusion (Pronadisa), supplemented with 0.1% cysteine, and 20% of a 200 g/l skim milk (Scharlab) solution. This mixture was then diluted 1:2 (v/v) in the same medium and stored in aliquots at −80 °C. Mice faecal pellets from the animal groups were collected before the experiments and preserved using the same procedure.

### Antibiotic treatment and microbiota stool transplant

Four groups of five C57BL/6J female 6 week-old mice were used in the present study. Three groups were treated with an antibiotic cocktail composed of 1 g/l ampicillin, 1 g/l metronidazole, 1 g/l neomycin and 0.5 g/l vancomycin in drinking water, as previous described with modifications [46]. To diminish intestinal microbiota load, the animals were given the antibiotic-containing water *ad libitum* for three weeks and the antibiotic cocktail was renewed every three days. To determine the number of viable bacteria in mice stools, samples were collected at the beginning of the experiment and the day before RV infection, and serially diluted in brain heart infusion supplemented with 0.1% cysteine. The dilutions were plated in Wilkins-Chalgren medium containing 1.5% agar and incubated in anaerobic conditions (AnaeroGen, Sigma) at 37 °C for 48 h. Mice were fed a standard diet until one week before fecal transplantation, when it was substituted by purified-defined germ-free diet (AIN-93G, Envigo).

The control group was maintained without antibiotics. Twenty-four hours after antibiotic treatment completion, two groups of mice were subjected to faecal material transplantation (FMT) as previously described with modifications [47]. One group of mice was transplanted with the preserved microbiota from the same group of mice before antibiotic treatment (100 μl of prepared faecal material per mice for three consecutive days through oral gavage). A second group of mice received a FMT using the same procedure with a pool of bacteria from infant faeces (100 μl of a pooled mix of four healthy infants for three days through oral gavage).

### Rotavirus challenge

Six days after FMT the mice were orally inoculated with 1×10^10^ genome copy equivalent (GCE) of RV_wa_ in 100 μl of TNC. After RV dosing, the antibiotic-treated group without FMT continued the drinking water antibiotic treatment, whereas in the FMT mice groups, antibiotics were omitted from water from the day before microbiota transplant. Stool samples were collected daily for 7 days and kept at −20°C. The mice were euthanized at 7 days post-infection (dpi) and the small intestine was removed and stored in RNAlater (Sigma) at −80 °C for further analysis.

### Quantification of rotavirus from stool samples by RT-qPCR

We extracted RNA from mice stool samples using the NucleoSpin^®^ RNA Virus (Macherey-Nigel) kit following the manufacturer’s instructions. The primers and probe sequences utilized for RT-qPCR have previously been described [45]. Amplification of RNA samples was performed with one-step TaqMan RT-qPCR using the RNA UltraSense One-Step quantitative system (Thermo Fisher Scientific). The standard curve was generated by serial end-point dilution, amplifying 10-fold dilutions of the quantified plasmid containing the RV target sequence by RT-qPCR in triplicates.

### Quantification of cytokine and glycosyltransferase mRNA expression level

One hundred milligrams of tissue from the small intestine of infected mice were homogenized in 1 ml of Trizol (Thermo Fisher Scientific) using a Polytron PT10-35 GT (Thermo Fisher Scientific) at 16,000 rpm. After tissue disruption the RNA was purified following the manufacturer’s instructions and the RNA finally resuspended in 50 μl of DEPC-treated water. The RNA was treated with RNAse-free DNAse I (Thermo Fisher Scientific) to remove the contaminant DNA and retro-transcribed to cDNA using the SuperScript III enzyme (Thermo Fisher Scientific) and random primers following the manufacturer’s instructions. The cDNA obtained was kept at −20 °C until use. Expression levels of genes encoding cytokines IL1β, IL4, IL6, CXCL15, IL10, IL12, IL13, TNFα, IFNγ and TLR2 were studied in a LightCycler480 Instrument SW1.5 (Roche Life Science) and the expression analysis performed with the Rest Software [48]. The expression level of *GAPDH* and *RPLPO* housekeeping genes was used as reference. A list of primers employed can be found in Supplementary Table 1.

Glycosyltransferase FUT2 gene expression was also analysed by RT-qPCR. Primer/probe preparation for the gene was purchased from Integrated DNA Technologies, Inc. (IDT, assay ID: Mm.PT.58.50508299). The NZYSpeedy qPCR Probe Master Mix (NZY) was utilized. The qPCR was run in the StepOnePlus Real-Time PCR System (Applied Biosystems) and expression analysis was performed with Rest Software [48], using *HPRT* gene expression (assay ID: Mm.PT.39a.22214828) as a reference.

### Microbiota profiling in mice stool samples

To assess microbiota composition, DNA was extracted from mice faecal pellets obtained prior to RV_wa_ inoculation from all experimental groups with the Master Pure™ kit (Epicentre), including a bead beater treatment with 0.5 g of 0.1 mm glass beads in the lysis step (FastPrep 24-5G Homogenizer; MP Biomedicals, CA, USA), followed by final purification of the extracted DNA samples with mi-PCR Purification kit (Metabion). DNA was quantified with a Qubit 2.0 fluorometer (Invitrogen). Bar-coded amplicons of the 16S rDNA V3–V4 region, multiplexed using Nextera XT Index Kit, were subject to 2 × 300 bp paired-end run in a MiSeq-Illumina platform (SCSIE, University of Valencia, Spain). Data were demultiplexed using Illumina bcl2fastq© program and reads were checked for quality, adapter trimmed and filtered using AfterQC and FastQC v0.11.8 (http://www.bioinformatics.babraham.ac.uk) tools. QIIME software V1.9.1 [49] was used to analyse tMiSeq sequencing data, including forward and reverse reads joining, chimera removal, data filtering and taxonomic annotation. Chimeric sequences were removed from the reads using the USEARCH 6.1 algorithm. Reads were clustered into operational taxonomic units (OTUs) based on a 97% identity threshold value. Sequence were aligned to the Greengenes core reference database (version 13.8) using PyNAST. Taxonomic assignment was made using the UCLUST classifier. A total of 4,356,240 non-chimeric reads were obtained, with a mean of 207,440 sequences per sample. Datasets were rarefied to the minimum library size (43,791 reads) and normalized by total-sum scaling prior analysis with Calypso 8.84 (http://cgenome.net/calypso/) and MicrobiomeAnalyst (https://www.microbiomeanalyst.ca/; [50] software pipelines. The raw sequencing fastq files were deposited in the SRA repository (http://www.ncbi.nlm.nih.gov/sra) under Bioproject ID PRJNA706108.

## Acknowledgements

This work was supported by Spanish Government (Ministerio de Economía y Competitividad) grants AGL2017-84165-C2-1-R to MJY, AGL2017-84165-C2-2-R to JRD. The study was also supported by Valencian Government grant IDIFEDER/2018/056 and by European FEDER founds. RGR is the recipient of a postdoctoral grant from the Valencian Government (APOST/2017/037). CSB is the recipient of a FPI predoctoral grant from the Spanish Government (RE2018-083315). SVV is the recipient of a predoctoral grant from the Valencian Government (ACIF/2016/437).

## Author Contributions

Conceptualization, JRD, JB, VM, RGR, ARC, MJY; Methodology, RGR, CSB, ARC, ARC, SVV, VM; Writing – Original Draft Preparation, JRD, VM; Review & Editing, ARC, JB, MJY, RGR, CSB; Project Administration JRD; Funding Acquisition, JRD, MJY.

## Conflicts of Interest

The authors declare no conflict of interest.

---

**Supplementary Figure 1.**
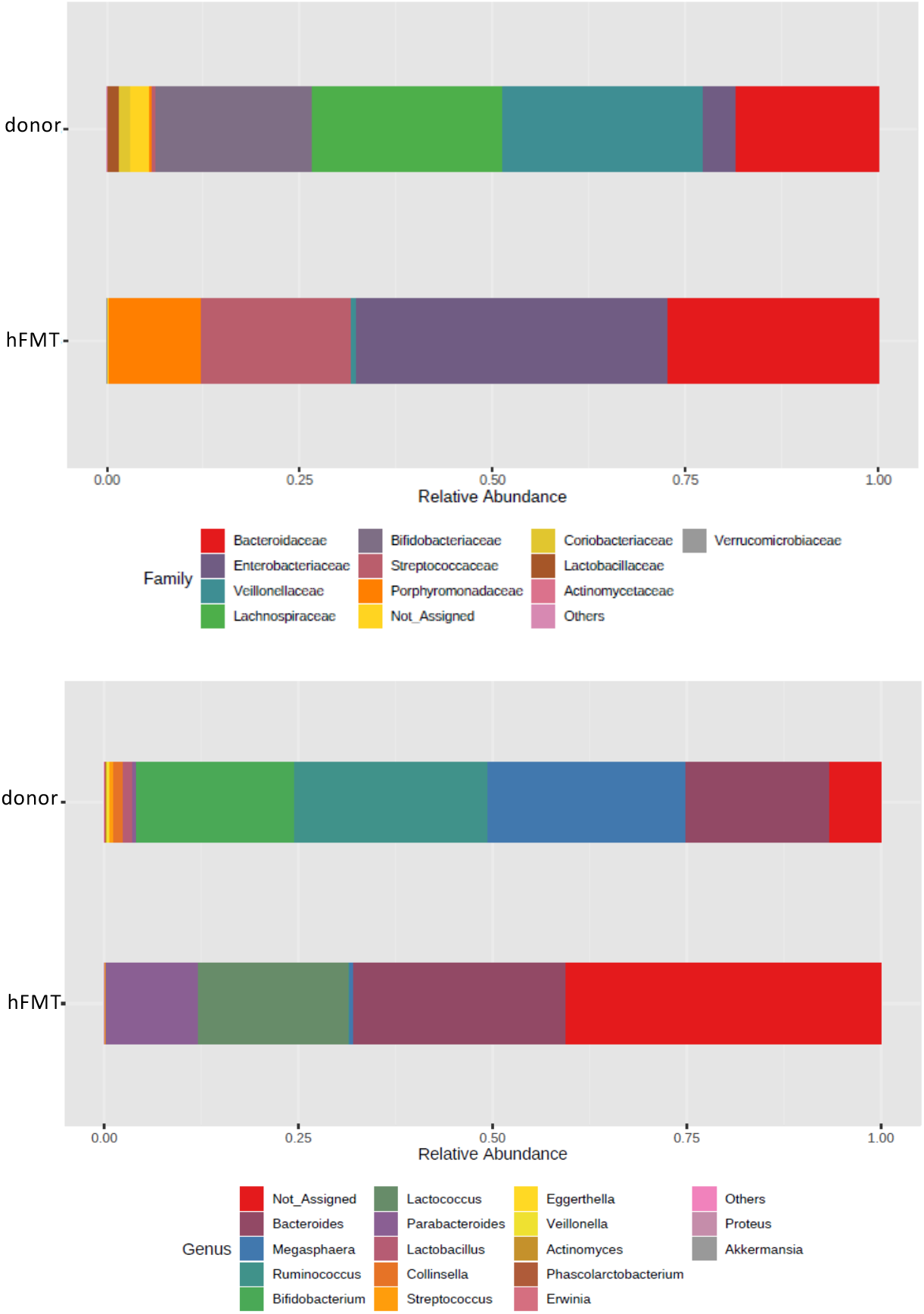
Relative abundances of bacterial taxa (family and genus) in donor faeces and faeces of mice subject to hFMT.

**Supplementary table 1.**
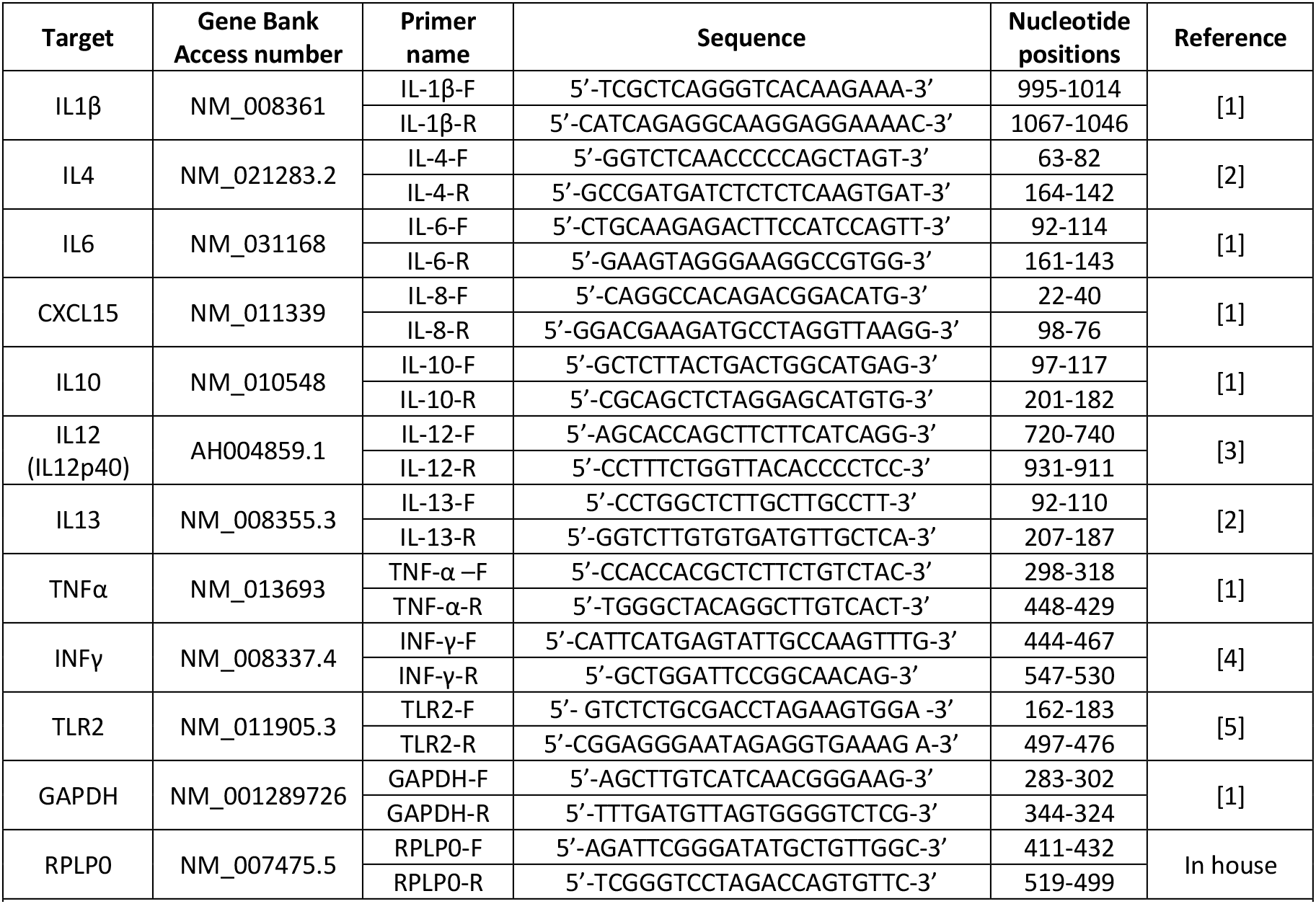

## Notes

### Competing Interest Statement

The authors have declared no competing interest.

